# metaExpertPro: a computational workflow for metaproteomics spectral library construction and data-independent acquisition mass spectrometry data analysis

**DOI:** 10.1101/2023.11.29.569331

**Authors:** Yingying Sun, Ziyuan Xing, Shuang Liang, Zelei Miao, Lai-bao Zhuo, Wenhao Jiang, Hui Zhao, Huanhuan Gao, Yuting Xie, Yan Zhou, Liang Yue, Xue Cai, Yu-ming Chen, Ju-Sheng Zheng, Tiannan Guo

**Author notes:** These authors contribute equally (Yingying Sun, Ziyuan Xing, Shuang Liang). Correspondence (Y. C.), (J. Z.), (T.G.).

## Abstract

**Background:** Analysis of mass spectrometry-based metaproteomic data, in particular large-scale data-independent acquisition MS (DIA-MS) data, remains a computational challenge. Here, we aim to develop a software tool for efficiently constructing spectral libraries and analyzing extensive datasets of DIA-based metaproteomics.

**Results:** We present a computational pipeline called metaExpertPro for metaproteomics data analysis. This pipeline encompasses spectral library generation using data-dependent acquisition MS (DDA-MS), protein identification and quantification using DIA-MS, functional and taxonomic annotation, as well as quantitative matrix generation for both microbiota and hosts. To enhance accessibility and ease of use, all modules and dependencies are encapsulated within a Docker container.

By integrating FragPipe and DIA-NN, metaExpertPro offers compatibility with both Orbitrap-based and PASEF-based DDA and DIA data. To evaluate the depth and accuracy of identification and quantification, we conducted extensive assessments using human fecal samples and benchmark tests. Performance tests conducted on human fecal samples demonstrated that metaExpertPro quantified an average of 45,000 peptides in a 60-minute diaPASEF injection. Notably, metaExpertPro outperformed three existing software tools by characterizing a higher number of peptides and proteins. Importantly, metaExpertPro maintained a low factual False Discovery Rate (FDR) of less than 5% for protein groups across four benchmark tests. Applying a filter of five peptides per genus, metaExpertPro achieved relatively high accuracy (F-score = 0.67–0.90) in genus diversity and demonstrated a high correlation (r_Spearman_ = 0.73–0.82) between the measured and true genus relative abundance in benchmark tests.

Additionally, the quantitative results at the protein, taxonomy, and function levels exhibited high reproducibility and consistency across the commonly adopted public human gut microbial protein databases IGC and UHGP. In a metaproteomic analysis of dyslipidemia patients, metaExpertPro revealed characteristic alterations in microbial functions and potential interactions between the microbiota and the host.

**Conclusions:** metaExpertPro presents a robust one-stop computational solution for constructing metaproteomics spectral libraries, analyzing DIA-MS data, and annotating taxonomic as well as functional data.

## Background

Microbial communities and functions have attracted increasing research interests in the past few years due to their crucial roles in human health, including nutrition, metabolism, and immunity^1^. Multi-omics approaches (*i.e.*, 16/18S ribosomal RNA sequencing, metagenomics) have been widely applied in gut microbiota studies to provide multifaceted information in characterizing the microbial profiles and their alterations linked with human diseases such as obesity, type 2 diabetes, hepatic steatosis, intestinal bowel diseases (IBDs), and cancer^2^. These technologies provide important information on the taxonomic composition and functional potential of microbiota but lack the messages of the truly expressed functions.

Metaproteomics is an emerging research area due to its unique strengths in quantifying the truly expressed proteins in the entire microbial community, assessing the community structure based on the biomass contributions of individual community members, exploring the interactions between microorganisms and their hosts or environment^3^, as well as identifying disease-associated protein biomarkers, *e.g.*, in human fecal^4^, or saliva^5^ samples.

However, data analysis of MS-based metaproteomics data is highly sophisticated. Searching against comprehensive protein databases containing several million protein sequences not only requires huge storage space and memory but also presents the tradeoff between proteome depth and false positive identifications^6^. Consequently, although widely used proteomics software tools like X!Tandem^7^, OMSSA^8^, MS-GF+^9^, Comet^10^, Proteome Discoverer (PD), and MaxQuant^11^ have been employed in metaproteomics data analysis, they are primarily applicable only to DDA-MS data. These tools are not well-suited for analyzing very large metaproteomic datasets (ranging from hundreds to thousands) due to suboptimal computational efficiency. Therefore, the majority of published metaproteomic datasets consist of fewer than 200 MS injections. To enhance computational efficiency, specialized software such as metaLab^12–14^, MetaProteome Analyzer (MPA)^15^, and ProteoStorm^16^ have been developed exclusively for metaproteomics analysis. However, they are all designed for DDA-MS-based metaproteomics analysis. Data-independent acquisition mass spectrometry (DIA-MS) demonstrates superb reproducibility, throughput, and proteome depth for single-injection analysis of complex proteomes^17^. However, DIA-MS generates highly convoluted fragment ion spectra which require sophisticated data analysis^18^, especially in metaproteomic samples that have an increased chance of co-elution of precursor ions^19^. Only two software tools namely diatools^20^ and its updated version glaDIAtor^21^ were designed for DIA-MS-based metaproteomics analysis.

However, neither of them is compatible with parallel accumulation-serial fragmentation combined with data-independent acquisition (diaPASEF) data which include ion mobility information^22^. In particular, diaPASEF achieves almost 100% peptide precursor ion current for DIA-MS data acquisition, leading to 5–10 times higher sensitivity improvement, but further increasing the complexity of metaproteomic data. Reducing search space without compromising proteomic depth is crucial for diaPASEF-based metaproteomics data analysis. Spectral library-based database search methods following peptide prefractionation typically yield a higher number of identified spectra compared to library-free database and pseudospectral library search methods^23^ in DIA analysis. Moreover, this approach requires less computational resource due to a reduced search space compared to library-free database search methods^21^. FragPipe^24^ harnesses the remarkable speed of the MSFragger proteomic search engine, surpassing X!Tandem, SEQUEST, and Comet by 100-fold in the analysis of a single DDA-MS run consisting of 41,820 MS/MS spectra. It seamlessly supports both Orbitrap and PASEF DDA-MS data. Additionally, FragPipe’s split database function, coupled with an accelerated proteinprophet module, renders it highly suitable for spectral library generation in metaproteomics data^25^. DIA-NN^26^ facilitates comprehensive proteome quantification in DIA-MS data, proving particularly advantageous for high-throughput applications owing to its rapid processing. Notably, a recent study by Demichev *et al*. demonstrated that integrating FragPipe with DIA-NN for diaPASEF data analysis led to a substantial increase in proteomic depth, approximately 70% higher than the originally published diaPASEF workflow using DIA-NN library-free analysis^27^.

Based on these progresses, here we developed a metaproteomic data analysis workflow called metaExpertPro, which is compatible with both DDA and DIA MS data from both ordinary MS and MS with ion mobility information such as timsTOF. metaExpertPro utilizes DDA-MS data for spectral library generation and DIA-MS data for peptide and protein identification and quantification. It offers a comprehensive one-stop metaproteomic workflow, including peptide and protein measurement, functional and taxonomic annotation, and quantitative data matrix generation. Additionally, metaExpertPro is easily accessible as a Docker image (https://github.com/guomics-lab/metaExpertPro). This method showed deep identification of about 45,000 peptides per human fecal sample from more than 10,000 protein groups with a 60 min LC gradient DIA-MS acquisition on a timsTOF Pro. Benchmark tests demonstrated that metaExpertPro maintains both low factual FDR (∼ 5%) and high-sensitivity identification at protein group level. Also, laboratory-artificial microbial mixture tests showed that metaExpertPro achieves high accuracy in both diversity and relative abundance at genus level. Furthermore, the negligible effects of different databases on quantification suggest that matched metagenomic sequencing is not required, and the results generated by metaExpertPro based on public different databases will be directly comparable. Finally, we applied the metaExpertPro software to study fecal specimens from dyslipidemia (DLP) patients. The results uncovered previously unclear alterations of microbial functions and the potential interactions between the microbiota and the host.

## Results

### Overview of metaExpertPro workflow

In this study, we proposed a metaproteomics data analysis workflow called metaExpertPro for the measurement of peptides, protein groups, functions, and taxa of gut microbes as well as host proteins based on DDA-MS and DIA-MS data from either Thermo Fisher Orbitrap (.raw / .mzML format) or Bruker (.d format) mass spectrometers. Briefly, the workflow includes four stages: DDA-MS-based spectral library generation, DIA-MS-based peptide and protein quantification, functional and taxonomic annotation, as well as quantitative matrix generation. The implementation of the metaExpertPro workflow is shown in Figure 1A with more details explained below.

**Figure 1.**
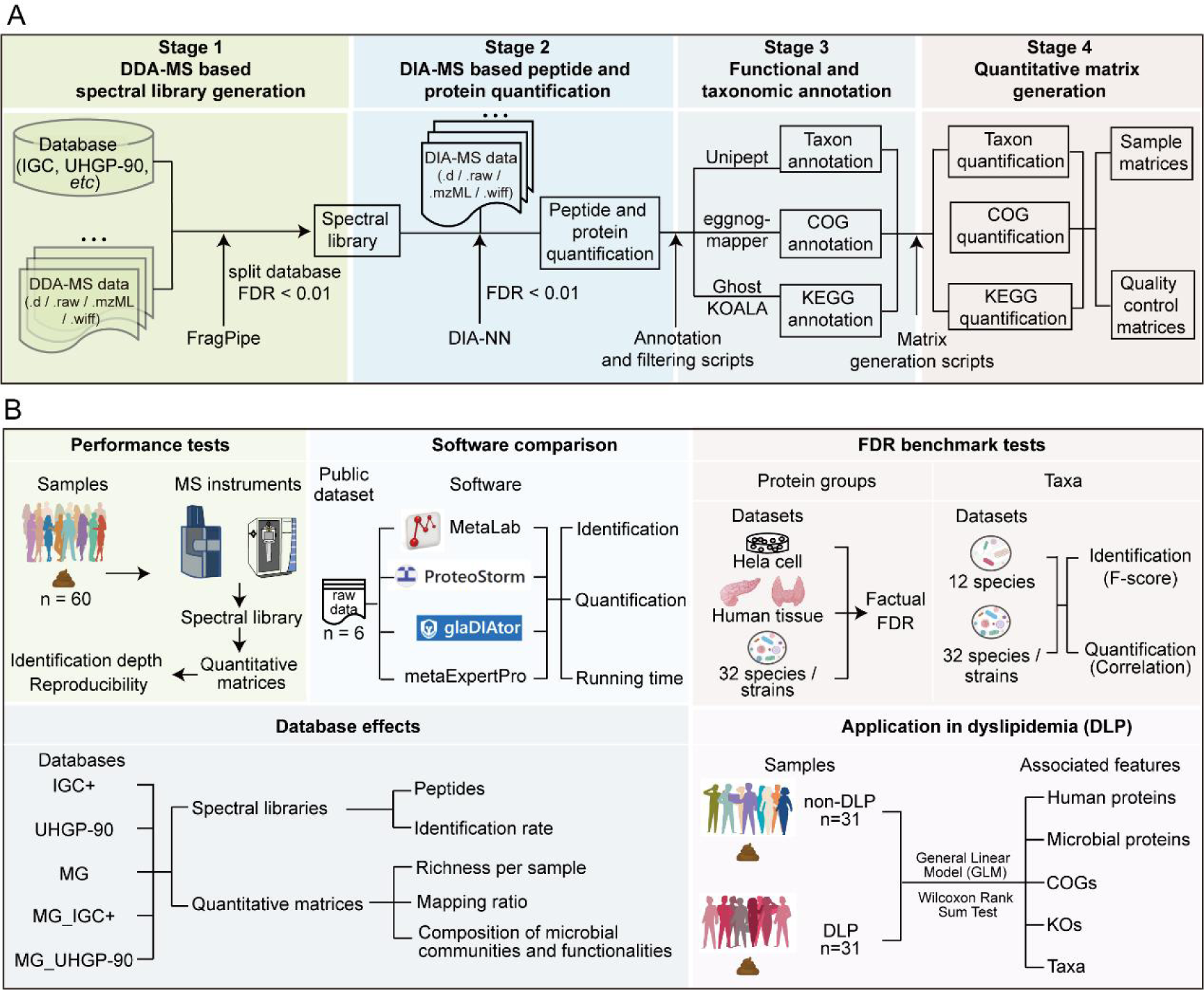
Overview of computational workflow and performance tests of metaExpertPro. (A) Overview of metaExpertPro workflow. The metaExpertPro workflow consists of four stages, including DDA-MS based spectral library generation, DIA-MS based peptide and protein quantification, functional and taxonomic annotation, as well as quantitative matrix generation. Stage 1 depicts the spectral library generation process using FragPipe software. Detailed procedures are described in Methods. DDA-MS raw data in .d, .raw, .mzML, .wiff formats are all compatible. In stage 2, the peptides and proteins are quantified based on DIA-MS data and the spectral library using DIA-NN. In stage 3, the taxa, COGs, and KEGGs are annotated by Unipept, eggnog-mapper, and GhostKOALA, respectively. The annotation results are then filtered through the in-house filtering scripts. In stage 4, the quantitative matrices of subject samples and quality control samples at taxa, COG, and KEGG levels are generated using matrix generation scripts. (B) Overview of performance tests of metaExpertPro. The identification depth and reproducibility of metaExpertPro were assessed in 60 human fecal samples, with MS raw data acquired using both timsTOF Pro and Orbitrap instruments. The results of identification and quantification, as well as running time were compared among MetaLab, ProteoStorm, glaDIAtor, and metaExpertPro software tools utilizing a public dataset. FDR benchmark tests were performed at both the protein groups and taxa levels using multiple datasets. At the protein group level, factual FDR was employed to gauge the accuracy of protein group identification. At the taxon level, the F-score was calculated for the identification accuracy test, while the correlation was computed for the quantification accuracy test. The impact of databases on spectral libraries and quantitative matrices was assessed using IGC+, UHGP_90, MG, MG_IGC, and MG_UHGP-90 databases. Finally, metaExpertPro was employed for metaproteomics data analysis on dyslipidemia (DLP) and non-DLP samples to characterize DLP-associated features at the human protein, microbial protein, COG, KO, and taxon levels.

In the first stage, we applied FragPipe (version 20.0) software^24^ for spectral library generation (Figure 1A). To minimize computational memory demands, the original database (*e.g.* integrated gene catalog database (IGC) of human gut microbiome and Unified Human Gastrointestinal Protein (UHGP)) was divided into multiple databases utilizing the database split parameter of MSFragger. The more the database is split, the less memory is required, but the longer the runtime. Therefore, users need to judiciously choose the number of database splits based on the quantity of protein sequences contained in the database. Then, each DDA-MS raw data was searched against each split database, generating a pepXML and a pin file. All the pepXML and pin files for each DDA-MS raw data were aggregated for PSM validation using either PeptideProphet or MSBooster-Percolator. To decide the appropriate PSM validation method, we assessed the number of protein groups and the factual FDR in two benchmark tests using PeptideProphet and the MSBooster-Percolator method, respectively. The benchmark tests utilized the public dataset (PXD006118) from a synthetic community of 32 organisms, searching against a sample-matched metagenomic database supplemented with either a subset of IGC database, containing ten times the proteins in metagenomic database, or 48 human gut microbial species. False positives included contaminant proteins, IGC proteins, or proteins from the added microbial species (Figure S1A). Both benchmark tests demonstrated a lower factual FDR using the PeptideProphet method (0.057 vs 0.091 and 0.037 vs 0.048), despite the MSBooster-Percolator method achieving 8.7–12.1% higher protein group identifications than the PeptideProphet method (Figure S1B). To maintain a relatively low factual FDR, we selected PeptideProphet as the default PSM validation method in metaExpertPro.

In the second stage, we applied DIA-NN software ^26^ to identify and quantify peptides and proteins from each DIA-MS data file (Figure 1A). In the third stage, we performed taxonomic annotation using the peptide-centric taxonomic annotation software Unipept^28,29^, which has been proved to exhibit more accurate and precise taxonomic annotation^30^ compared to Kraken2^31,32^ and Diamond^33,34^. Because the Unipept only indexes perfectly cleaved tryptic peptides^35^, we *in silico* digested the peptides and filtered the peptide length before the Unipept taxonomic annotation (Figure S1B). To enhance annotation confidence, peptides with conflicting taxon annotations were excluded (Figure S1D). To eliminate unreliable taxa, we calculated the number of peptides associated with each taxon and selected taxa with more than 1, 3, 5, 10, 15, and 20 peptides. The metaproteomic functional annotation tools eggnog-mapper^36,37^ and GhostKOALA^38^ were integrated into the pipeline to process functional annotation (Figure 1A).

In the fourth stage, we generated quantitative matrices at nine levels including human peptide, microbial peptide, human protein group, microbial protein group, COG, KO, COG category, KO category, and taxonomy. The peptides corresponding to both human protein group and microbial protein group were removed from the quantitative results to avoid protein assignment ambiguity (Figure S2).

In summary, the metaExpertPro pipeline integrates high-performance proteomic analysis tools—FragPipe and DIA-NN—along with functional and taxonomic annotation software tools, employing rigorous filter criteria to provide a comprehensive metaproteomics workflow in a single package.

Subsequently, to assess the performance of metaExpertPro in human gut microbial samples, we conducted tests to evaluate identification depth and result reproducibility using two MS instruments. Additionally, we compared the measurement results and runtime of metaExpertPro with three existing metaproteomics software tools—MetaLab, ProteoStorm, and glaDIAtor. For workflow accuracy estimation, we computed the factual FDR of protein groups, the F-score of taxa, and the correlation between measured taxa and true protein amounts in multiple benchmark tests. Furthermore, we examined the effects of databases on spectral libraries and quantitative matrices using five mainstream human gut microbial databases. Finally, we applied metaExpertPro in metaproteomic analysis of dyslipidemia patients to explore potential associations between human gut microbial functions and taxa related to dyslipidemia (Figure 1B). Detailed descriptions of all tests are provided below.

### In-depth identification and high reproducibility of metaExpertPro workflow in human fecal samples

To demonstrate the benefits of metaExpertPro, we applied it to the metaproteomic analysis of 62 human fecal samples from 62 middle-aged and elderly volunteers of the Guangzhou Nutrition and Health Study (GNHS)^39^. Sixty samples were acquired using two MS instrument platforms: the timsTOF Pro (Bruker) and the Orbitrap Exploris™ 480 (Thermo Fisher Scientific) (Figure 2A). Approximately 5 μg peptides from each sample were mixed into a pooled sample for high-pH fractionation. A total of 30 fractionated samples were obtained. Each fraction was analyzed by DDA-MS acquisition with a 60 min gradient for spectral library generation. The remaining peptides from each sample were used for DIA-MS acquisition (Figure 2A). A total of 220,365 peptides and 58,952 protein groups, including 57,862 microbial protein groups and 1,065 human protein groups, were identified in the spectral library derived from timsTOF Pro (Figure 2 B). Using Exploris 480, 189,808 peptides and 51,269 protein groups, including 50,218 microbial protein groups and 1,024 human protein groups, were characterized (Figure 2C). The average identification rate of the acquired MS spectra was 32.2% and 29.3% for the spectral libraries derived from timsTOF Pro and Exploris 480, respectively (Figure 2 D–E, Table S1). The identification rates were comparable to the MetaPro-IQ^12^ results (medium = 32%) obtained from 4 h gradient DDA-MS run on the Q Exactive MS spectrometer for eight human stool samples. For each sample, we quantified 43,194 ± 11,704 (mean ± SD) microbial peptides corresponding to 15,501 ± 3,880 microbial protein groups, and 2,453 ± 398 human peptides corresponding to 537 ± 91 human protein groups on timsTOF Pro. On Exploris 480, we quantified 22,460 ± 4,964 microbial peptides corresponding to 11,301 ± 2,172 microbial protein groups, and 1,374 ± 246 human peptides corresponding to 414 ± 69 human protein groups (Figure 2 F–G, Table S2). Nevertheless, to the best of our knowledge, the numbers of peptide identifications on two types of MS instruments are the highest compared to the published metaproteomic results with the same or even longer LC gradient. For example, the MetaPro-IQ workflow identified 15,210 peptides per human fecal sample with 4 h gradient DDA-MS acquisition^12^, and glaDIAtor identified 8211 peptides per human fecal sample with 90 min gradient DIA-MS acquisition^21^. Moreover, the number of peptides with 60 min gradient acquisition on timsTOF Pro identified by metaExpertPro is comparable to the MetaPro-IQ results with 22 h of MS analysis (45,647 vs 44,955 peptides per human fecal sample)^40^. Due to the in-depth identification of peptides and protein groups, we also quantified an average of 90–92 microbial species, 68–71 genera, 1,406–1,511 COGs, and 1,350–1,475 KOs per human fecal sample (Figure 2 F–G, Table S2).

**Figure 2.**
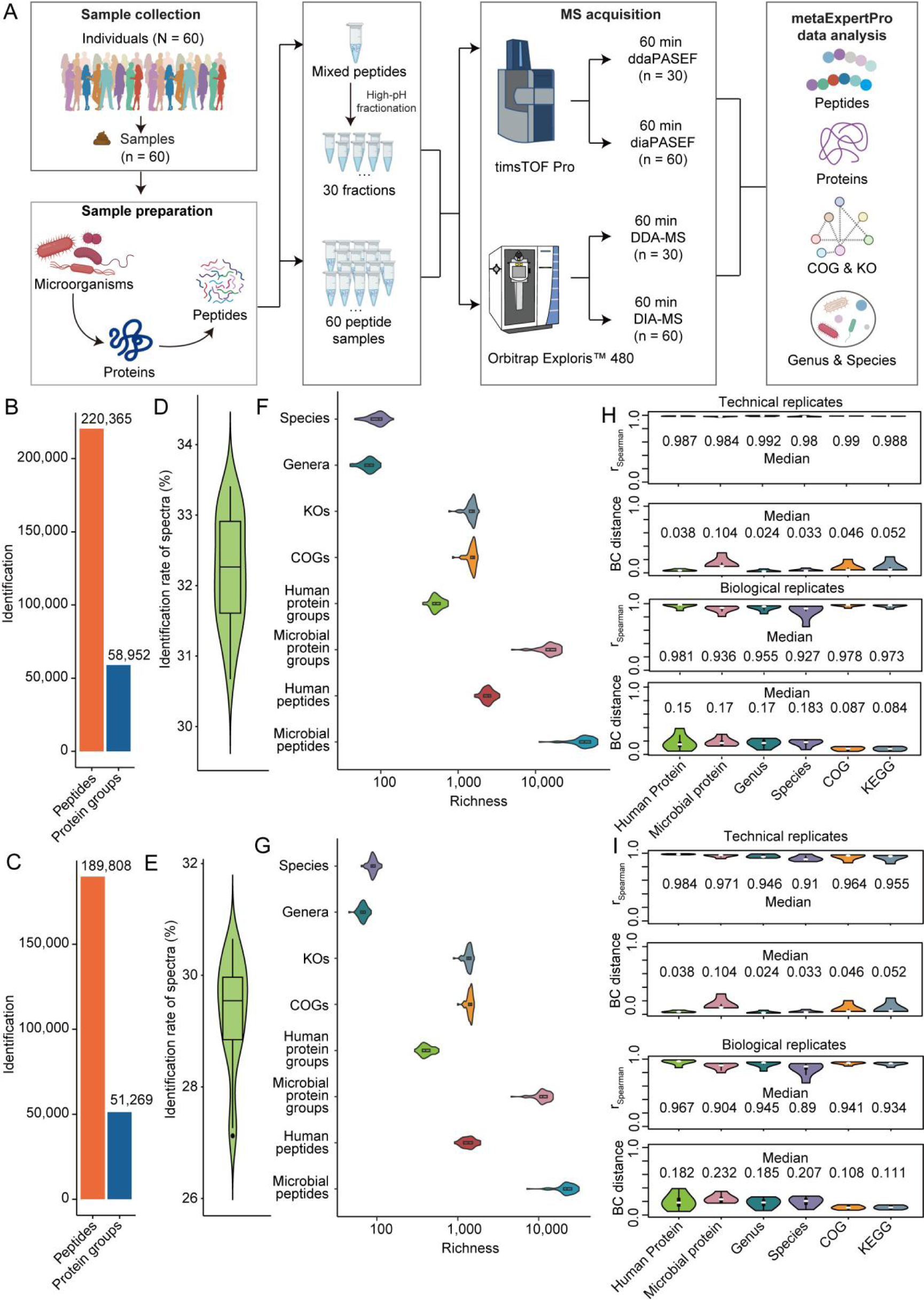
In-depth identification and high reproducibility of the metaExpertPro workflow in the metaproteomic analysis of human fecal samples. (A) Experimental design including sample collection, sample preparation, MS acquisition, and metaExpertPro data analysis of human fecal samples. A total of 60 peptide samples were obtained from 60 human fecal samples after the sample preparation process. For the DDA-MS based spectral library generation, the 60 peptide samples were firstly mixed. Then, the mixed peptides were fractionated into 30 fractions for DDA-MS acquisition. For DIA-MS based peptide and protein quantification, all 60 peptide samples were used for DIA-MS acquisition. Two types of mass spectrometers including timsTOF Pro and Orbitrap Exploris™ 480 were applied for both DDA-MS and DIA-MS acquisition. (B–C) Identification performance of peptides and protein groups in spectral libraries based on 30 DDA-MS runs on timsTOF Pro (B) or Orbitrap Exploris™ 480 (C) MS spectrometer. (D–E) Identification rate of the MS spectra acquired from 60 DIA-MS runs collected on timsTOF Pro (D) or Orbitrap Exploris™ 480 (E) MS spectrometer. The y-axis stands for the identification rate of acquired MS spectra (%). (F–G) The richness per sample detected on timsTOF Pro (F) or Orbitrap Exploris™ 480 (G) instrument. The x-axis reports the richness per sample at each level. (H–I) Pairwise Spearman correlation and Bray-Curtis (BC) distance between five pairs of technical replicates and six pairs of biological replicates based on timsTOF Pro (H) or Orbitrap Exploris™ 480 (I) instrument.

Another major benefit of DIA methods is the high degree of quantitative consistency. Thus, we next investigated the reproducibility of the quantified protein groups, functions, and taxa in five pairs of technical replicate samples and six pairs of biological replicate samples. As expected, high correlation was observed in all pairs of technical replicates at each level in two MS instruments (Figure 2 H–I). We also observed high correlation in all pairs of biological replicates at each level (Figure 2 H–I). In addition, the Bray-Curtis (BC) distance between all pairs of technical and biological replicates was low, and no statistically significant difference were observed between the first and the second repeat MS acquisition (PERMANOVA *p* = 0.89–1) (Figure 2 H–I).

The reproducibility between two MS instruments was assessed by comparing their identifications in the DDA-MS-based spectral library. Among the total peptides identified, 34.2% (104,521) were detected by both MS instruments, while 37.8% (30,291) of the total protein groups were identified by both instruments. These shared identifications accounted for 55.0% of the peptides and 58.9% of the protein groups identified by the Exploris 480 MS instrument (Figure S3A). For the DIA-MS-based quantification, 25.6% (56,939) of the total peptides and 36.2% (22,597) of the total protein groups were quantified by both MS instruments. The abundance correlation between the datasets generated by the two MS instruments were assessed using twelve biological replicate samples. The results showed that the median Spearman correlation was 0.788 for human proteins, 0.604 for microbial proteins, 0.673 for human peptides, 0.643 for microbial peptides, 0.908 for genera, 0.861 for species, 0.880 for COGs, and 0.852 for KOs, respectively (Figure S3B). In summary, metaExpertPro offers comprehensive identification and quantification capability for metaproteomics analysis of human fecal samples, utilizing MS raw data from either timsTOF or Exploris 480 instruments. Notably, it demonstrates remarkable reproducibility across replicate samples and MS instruments, ensuring reliable and consistent results.

### Comparison of metaExpertPro with other metaproteomics software tools

We next compared the application scenarios and the performance of metaExpertPro with the existing metaproteomics software tools. Among them, metaLab^13^, MetaProteomeAnalyzer (MPA)^15,41^, and ProteoStorm^16^ are DDA-MS-based metaproteomics analysis tools. They are all compatible with Q Exactive and Orbitrap Exploris MS instruments. Additionally, ProteoStorm is also compatible with Low-res LCQ/LTQ (Figure 3A). Both metaLab and MPA can perform DDA-MS-based peptide and protein quantification in metaproteomics analysis. Furthermore, metaLab provides additional functionalities for function and taxonomic annotation, as well as quantification. glaDIAtor^21^ is the next generation of diatools^20^. diatools and glaDIAtor are currently the only published analysis tools available for DIA-MS metaproteomics. However, it is important to note that neither glaDIAtor nor diatools is compatible with PASEF MS instrument. metaExpertPro is the exclusive DDA-assisted DIA-based metaproteomics analysis tool that is compatible with the timsTOF MS instrument. It provides a comprehensive solution encompassing DDA-MS-based spectral library generation, DIA-MS-based peptide and protein quantification, as well as function and taxonomic annotation and quantification, all in one platform (Figure 3A). To compare the performance of these software tools, we reanalyzed the Orbitrap acquired DDA-MS and DIA-MS datasets from six human fecal samples published by the Elo team^42^. For the DDA-MS-based software tools metaLab and ProteoStorm, six DDA-MS raw data files were used for peptide and protein quantification or identification. On the other hand, in the case of DIA-MS-based software tools glaDIAtor and metaExpertPro, these same six DDA-MS data sets were employed for spectral library generation. Subsequently, peptide and protein quantification were performed using DIA-MS raw data and the generated spectral library (Figure 3B).

**Figure 3.**
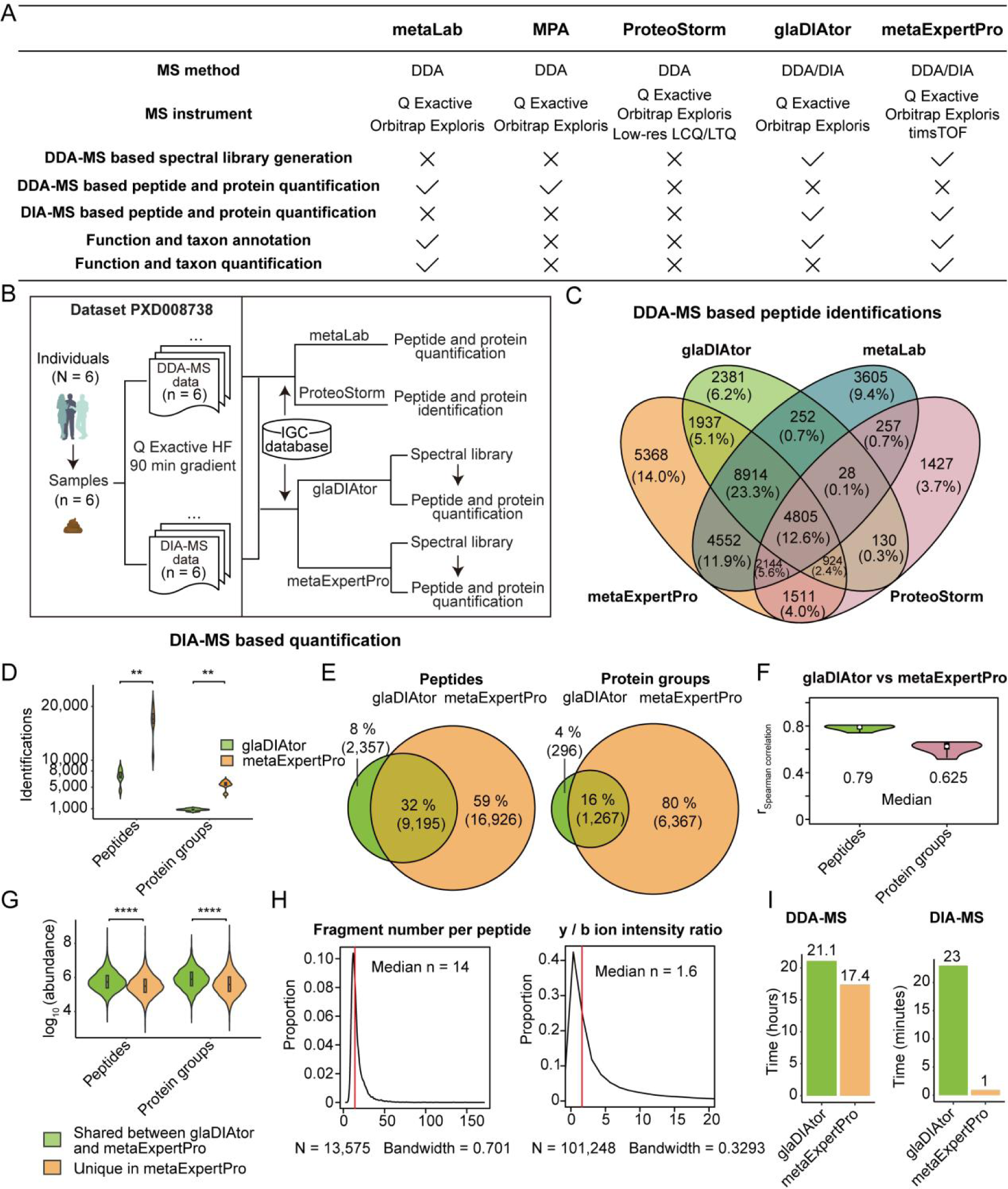
Comparison of metaExpertPro with other metaproteomics software tools. (A) Comparison of the application situations among metaLab, MetaProteomeAnalyzer (MPA), ProteoStorm, glaDIAtor and metaExpertPro. (B) Experimental design of the comparison between two software packages. The DDA-MS and DIA-MS data from dataset PXD008738 and the integrated gene catalog database (IGC) database were used for the measurement of peptides and protein groups by the metaLab, ProteoStorm, glaDIAtor, or metaExpertPro. (C) Comparison of peptide identifications by metaLab, ProteoStorm, glaDIAtor, and metaExpertPro. (D) The number of peptides and protein groups quantified by glaDIAtor and metaExpertPro. (E) The overlapped peptides or protein groups quantified by glaDIAtor and metaExpertPro. (F) The Spearman correlation of the abundance of peptides and protein groups quantified by both glaDIAtor and metaExpertPro. (G) Comparison of the intensity of peptides or protein groups identified by both glaDIAtor and metaExpertPro or identified by metaExpertPro only. (H) Density plots of the fragment number and the *y / b* ion intensity ratio of each peptide. The red line shows the median of the fragment number per peptide or the *y / b* ion intensity ratio. (I) Comparison of the running time between gladiator and metaExpertPro in DDA-MS based spectral library generation and DIA-MS based quantification. The tests were performed using six DDA-MS and six DIA-MS raw data of human fecal samples in dataset PXD008738 on AMD EPYC hardware and a 512G RAM computer. *p* value: * *p* < 0.05; ** *p* < 0.01; *** *p* < 0.001, **** *p* < 0.0001.

We compared DDA-MS-based peptide identifications among metaExpertPro, glaDIAtor, metaLab, and ProteoStorm. metaExpertPro demonstrated the highest peptide identifications in the spectral library (30,155) among the compared tools, surpassing glaDIAtor (19,371 peptides), metaLab (24,557 peptides), and ProteoStorm (11,226 peptides) in the spectral library. Despite the variations in peptide identification counts, metaExpertPro exhibited substantial overlap with other tools. It identified 16,580 peptides shared with glaDIAtor, 20,415 peptides shared with metaLab, and 9,384 peptides shared with ProteoStorm. These shared peptides accounted for 85.6%, 83.1%, and 83.6% of the total peptides identified by glaDIAtor, metaLab, and ProteoStorm, respectively (Figure 3C, Table S3). Additionally, metaExpertPro identified 5,368 unique peptides in the spectral library. Next, we compared the DIA-MS-based quantification of metaExpertPro and glaDIAtor. To ensure a fair comparison, both software tools were set to DDA-assisted DIA mode, guaranteeing identical raw data input for the analysis. Using metaExpertPro, we measured more than two-fold peptides (mean ± SD = 16,971 ± 3,315 vs 6,918 ± 1,456) and six-fold protein groups (mean ± SD = 5,368 ± 885 vs 812 ± 218) compared to glaDIAtor (Figure 3D, Table S4). Over half of all the peptides (59%) and protein groups (80%) were only detected by metaExpertPro. 32% of the peptides and 16% of the protein groups were quantified by both workflows. Only 8% of the peptides and 4% of the protein groups were quantified by glaDIAtor only (Figure 3E). In the comparison of peptide and protein abundance between the two workflows, we observed a relatively high correlation in the abundance of peptides and protein groups quantified by both metaExpertPro and glaDIAtor (median r_Spearman_ = 0.79 and 0.63) (Figure 3F). Furthermore, the abundance of peptides and protein groups exclusively detected by metaExpertPro was significantly lower compared to those identified by both workflows (Figure 3G, Table S5). These findings suggest that our workflow excels in identifying low-abundance peptides and protein groups.

To further verify the confidence of the peptides quantified only by metaExpertPro compared to glaDIAtor, we inspected the probability, the number of fragments, the *b / y* ion intensity ratio, and the spectra of these peptides. Among the 30,155 peptides identified in the metaExpertPro spectral library, 13,575 peptides were uniquely identified compared to glaDIAtor spectral library, while 16,580 peptides were shared between the two libraries. We firstly evaluated the accuracy of the 13,575 peptides in the metaExpertPro spectral library, confirming their reliability. Remarkably, all these peptides exhibited peptide probability values of 0.9963 ± 0.12 (median ± SD), indicating the high confidence in the peptide-spectrum matches. The median number of fragments matched for all peptides was 14, ranging from a minimum of 5 fragments to a maximum of 169 (Figure 3H). Notably, among the 13,575 peptides, 99.6% displayed two-sided fragment types, while only 54 peptides were identified as one-sided. Furthermore, in ion trap mass spectrometry, the intensities of *y*-ions are typically approximately twice that of their corresponding *b*-ions^43^. Among the 13,575 identified peptides, the median ratio of intensities between *y*-ions and their corresponding *b*-ions was 1.6, aligning with the anticipated pattern for complex peptide spectra (Figure 3H). To visually showcase the qualitative accuracy of the peptide identifications in the metaExpertPro spectral library, we obtained the DDA MS/MS spectra of the top 20 lowest abundant peptides. All 20 peptide spectra can be identified with at least 8 fragments containing both *y* ions and *b* ions. Most of the high-intensity peaks in the spectra can be matched to fragments, and there was a large dynamic range between high and low-intensity fragments. In addition, the intensity of *y* ions is higher than that of *b* ions (Figure S4). These criteria are in line with the manual assessment of high-quality peptide segments^43^, which demonstrate the reliability and precision of the identified peptides in the spectral library (Figure S4). Collectively, these findings strongly support the high quality and reliability of the peptides exhibiting relatively low abundance.

Next, we conducted a comparison of the running times for metaExpertPro and glaDIAtor on an AMD EPYC hardware system with 512 GB RAM using the PXD008738 dataset. With ten threads, glaDIAtor took approximately 21.1 hours for DDA-MS analysis, while metaExpertPro required approximately 17.4 hours. For DIA-MS analysis, glaDIAtor took around 23 minutes per file, while metaExpertPro completed the analysis in just 1 minute per file (Figure 3I).

Considering that the number of DDA-MS raw data is usually less than 100, while a high-throughput project may involve thousands of DIA-MS raw data files, metaExpertPro proves to be well-suited for high-throughput metaproteomic analysis.

In conclusion, the metaExpertPro workflow effectively enhanced proteome depth and upheld strong quantitative reproducibility in metaproteomic analysis. While the generation of DDA-MS-based spectral libraries using metaExpertPro may require longer running times, the DIA-MS-based quantification process is notably faster. This characteristic offers a significant advantage, particularly in high-throughput studies utilizing DIA-MS.

### Benchmark test of protein group identifications of metaExpertPro

We further investigated the accuracy of protein groups identified by metaExpertPro using benchmark tests. We initially assessed the factual FDR of protein groups in the spectral library using the published dataset of HeLa cells^30^. Briefly, the DDA-MS data of the HeLa cell was searched against the human protein database (Swiss-Prot, date 20211213) supplemented with 0×, 1×, 10×, 100×, and the entire mouse microbiome catalog sequences (∼2.6 million proteins), respectively (Figure 4A). The factual FDR is defined as the bacterial and contaminant hits divided by all the identified hits. As expected, when searching against the human protein database only (benchmark standard), the factual FDR was extremely low (0.015) (Figure 4B, Table S5). The count of human protein groups reached 5,511 (Figure 4B, Table S6), surpassing the originally published result of approximately 5,000 human protein groups^30^ based on a single-step search using MaxQuant software^11^. When increasing microbial sequences in the human protein database, the factual FDRs remained well-controlled (FDRs = 0.022–0.028), and the count of true human protein groups showed a slight decrease compared to the benchmark result (5,082 in the supplemented all bacteria sequences vs. 5,431 in the human protein database only) (Figure 4B, Table S6). To evaluate the ability of metaExpertPro to maintain a low protein-level FDR with larger sample sizes, we extended the number of DDA-MS raw data to 255, including 100 pancreas tissue samples and 155 thyroid tissue samples (IPX0001400000). These raw data were then searched against the human protein database (Swiss-Prot, date 20211213) supplemented with 0×, 1×, 10×, and 100× mouse microbiome catalog sequences (Figure S5A). The factual protein group FDRs remained below 5% when adding 0×, 1×, or 10× mouse protein sequences (∼2.6 million proteins) (Figure S5B, Table S7). However, when searching against 100× mouse protein sequences, the protein group FDR reached 5.4%. This suggests that controlling the factual protein group FDR becomes challenging when both the sample size and the unmatched protein sequences in the database increase in metaExpertPro.

**Figure 4.**
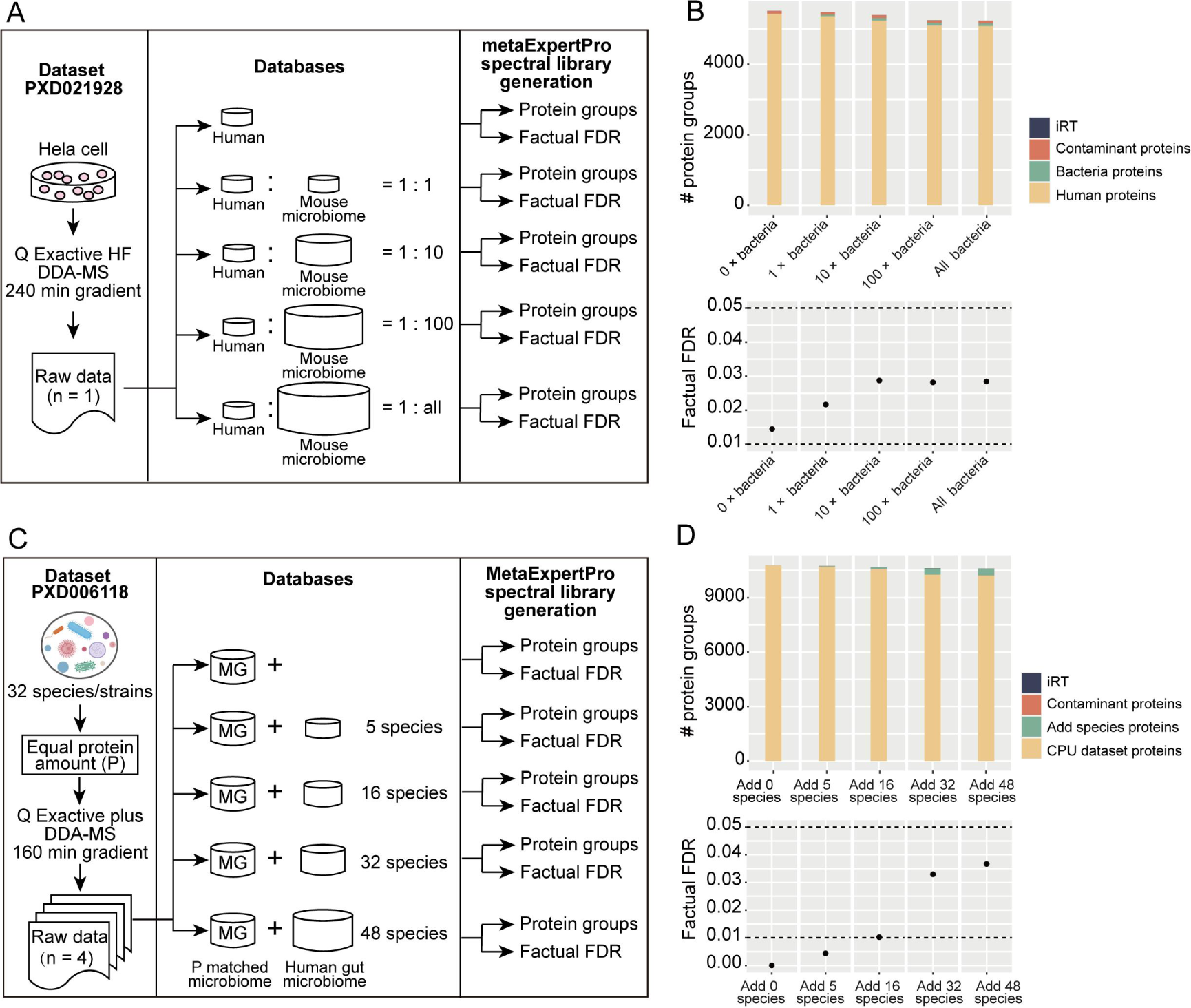
Benchmark test of protein group identifications of metaExpertPro. (A) The experimental design of the benchmark test for protein group identification based on Hela cell sample. The DDA-MS data of Hela cell from dataset PXD021928, and the databases containing human proteins supplemented with different sizes of mouse microbiome catalog were used for spectral library generation using metaExpertPro. (B) The number and factual FDR of the protein groups identified from HeLa cell MS raw data searching against the human protein database supplemented with 0×, 1×, 10×, 100×, and all the mouse microbiome catalog (∼2.6 million proteins), respectively. (C) The experimental design of the benchmark test for protein group identification based on bacteria mixture samples. The DDA-MS data of 32-species mixture from dataset PXD006118 (P: equal protein amount) were searched against databases containing P matched metagenomic database supplemented with 0, 5, 16, 32, and 48 human gut microbial species databases, respectively. (D) The number and factual FDR of protein groups in each subset test is present. The dashed lines depict the factual FDR of 0.05 and 0.01.

To gain insights into real-life scenarios of metaproteomics studies, we conducted two additional benchmark tests to identify false positive microbial proteins from microbiota mixtures. In the first test, we used the “equal protein amount” (P) dataset (PXD006118) and searched it against a metagenomic database (MG) supplemented with varying numbers of human gut microbiota species protein databases (5, 16, 32, 48) (Figure 4C). In the second test, we added the protein sequences of 0×, 1×, 5×, 10× IGC+ protein sequences (10,352,085) to the MG database (Figure S5C). Remarkably, we consistently achieved factual protein group FDRs below 5%, except for the 10× IGC+ benchmark test, which had a factual FDR of 5.8% (Figure 4D, Figure S5D, Table S8–S9). These results indicate the robustness of metaExpertPro in maintaining a low protein-level FDR in challenging scenarios.

In conclusion, the metaExpertPro workflow effectively maintains both a low factual FDR and high-sensitivity identification at the protein group level during spectral library building.

### Taxonomic accuracy estimation of metaExpertPro

Determination of taxonomic annotation and biomass contributions is another challenge due to a large number of homologous protein or peptide sequences derived from hundreds of closely related species. Thus, we next estimated the taxonomic accuracy at genus and species levels using two artificial bacterial community datasets. One of the datasets is the mixture of twelve different bacterial strains isolated from fecal samples of three human donors (hereafter referred to as “12-mix data”) published by Pietilä et al.^21^ (Figure 5A). Another dataset is called “CPU data” which were generated from synthetic communities consisting of 32 organisms with “equal cell number” (C), “equal protein amount” (P), and “uneven” (U) published by Kleiner and colleagues^44^ (Figure 5B). We searched the 12-mix data against the integrated gene catalog database (IGC) of human gut microbiome^45^ and the CPU MS data against the matched metagenomic database^44^ using the metaExpertPro workflow. Then, we calculated the true positive (TP), false positive (FP), false negative (FN), and F-score^46^ (the harmonic mean of precision and recall) at genus and species levels. When filtering out the taxa annotated by only one peptide, we got a relatively high true positive rate (TPR) (8/10) and a low false negative rate (FNR) (2/10) at genus level using the 12-mix dataset. But we also obtained a high false discovery rate (FDR) (10/18–11/19) and thus a relatively low F-score (average of 0.56) at genus level (Table S10). At species level, because of the decrease of TPR and increase of FNR and FDR, the F-score further decreased to 0.26 (Table S10). The average F-scores of the CPU data were 0.73 and 0.40 at genus and species level, respectively, outperforming the 12-mix data. Interestingly, the numbers of FP taxa in “uneven” samples were extremely low (4–5), resulting in high F-scores (0.84–0.86) at genus level (Table S10).

**Figure 5.**
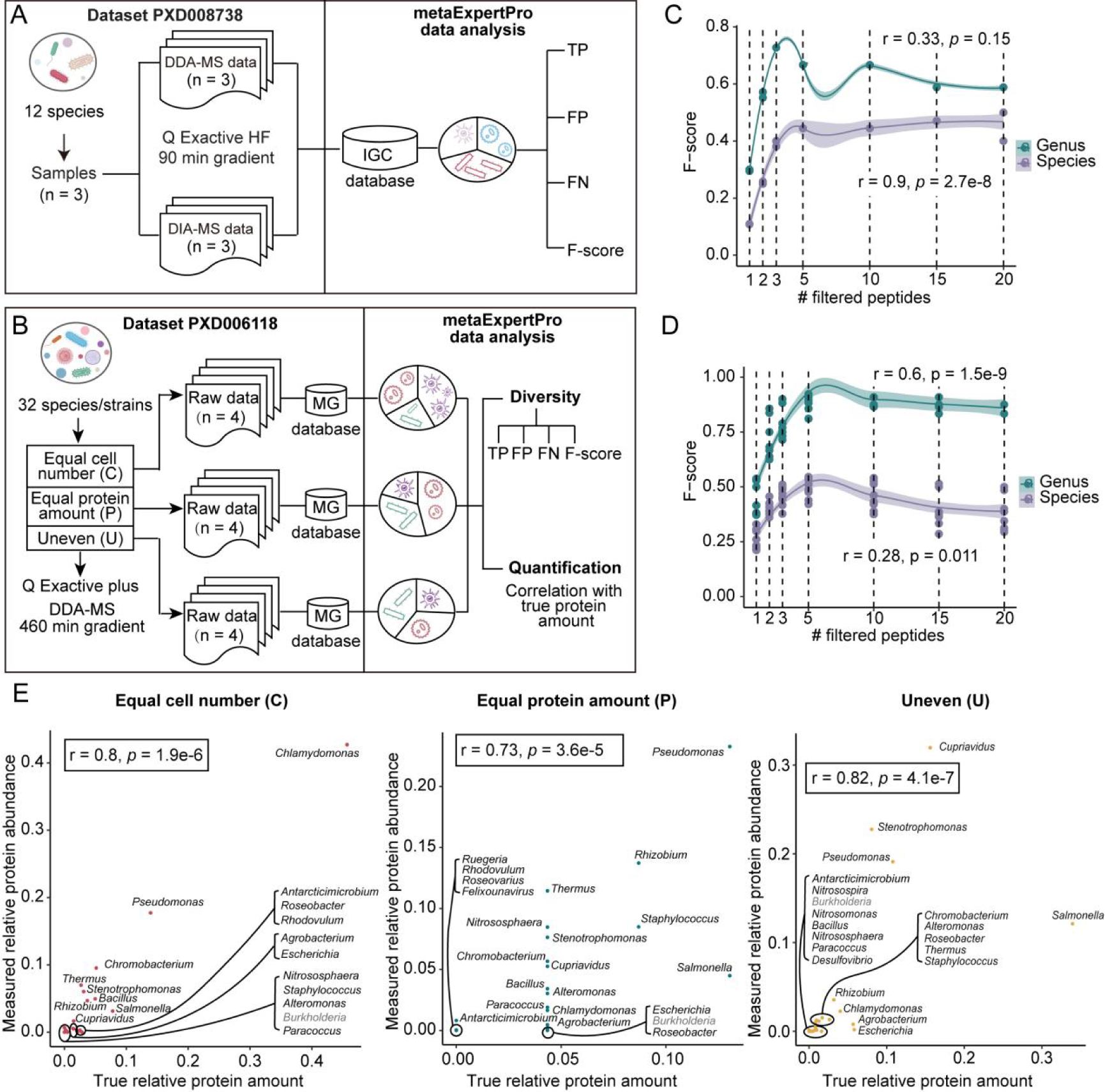
Taxonomic accuracy estimation of metaExpertPro. (A–B) The experimental design of the benchmark tests for taxa based on the 12-mix dataset (PXD008738) (A) and CPU dataset (C: equal cell number; P: equal protein amount; U: uneven) (PXD006118) (B). The figure depicts the original samples, the MS instrument, MS gradient, and MS acquisition modes applied in the 12-mix dataset (A) and CPU dataset (B). The 12-mix MS data were searched against the integrated gene catalog (IGC) database, while the CPU data were searched against the matched metagenomic database. The true positive (TP), false positive (FP), false negative (FN), and F-score of genera and species in each sample were calculated for both datasets. The measured relative abundance of genera or species was correlated with the true protein amount in the CPU dataset (B). (C–D) F-score of genera or species filtered by different numbers of corresponded peptides based on 12-mix MS data (C) or CPU MS data (D) using metaExpertPro. The F-score is the harmonic mean of precision and recall. The x-axis represents the minimum number of distinct peptides corresponding to genera or species. The y-axis displays the F-score of genera or species corresponding to the peptide count cutoff. The lines are smoothed by LOESS regression. (E) Spearman correlation between the true relative protein amount and the relative protein abundance of genera measured by metaExpertPro in the CPU dataset. The genera were filtered by containing at least five distinct peptides. Genera shown in grey indicate their absence in the Unipept database.

Here the F-scores were relatively low. Thus, we next investigated the impacts of the spectral count of peptides, the peptide length, and the number of peptides corresponding to the taxa on the TP and FP identifications at both genus and species levels (Figure S6 A–B). The data showed that, while all these three factors exhibited significant differences between the TP and FP identifications, the number of peptides corresponding to the taxa displayed the highest difference (Figure S6 A–B). After checking the peptide count distribution of TP and FP taxa (Figure S6 C–D), we filtered the number of peptides corresponding to taxa at the threshold of 1, 2, 3, 5, 10, 15, and 20, respectively, and recalculated the TF, FP, FN, and F-score. The data showed that filtering the taxa with at least five peptides led to the highest F-scores (C: 0.90; P: 0.85; U: 0.90) at the genus level (Figure 5 C–D, Table S10) in C, P, U datasets. This resulted in high TPR (C: 15/17; P: 15/17; U:17.25/20), low FNR (C:2/17; P: 2/17; U: 2.75/20) and low FDR (C: 1.5/16.5; P: 3.5/18.5; U:1/18.25). However, in the 12-mix dataset, filtering at least three peptides led to the highest F-scores (0.73) at the genus level. At the species level, we also obtained the highest F-score with the threshold of five peptides. But at the species level, the F-scores were still relatively low in two datasets (0.44–0.55) (Figure S6 E–F, Table S10).

The true quantitative information of the microorganisms in the CPU dataset^44^ allowed us to investigate the accuracy of the relative abundance of the taxa calculated by metaExpertPro workflow. With a threshold of five peptides, relatively high correlation between the true protein biomass of genera and the metaExpertPro results were observed (r_Spearman_ = 0.8, 0.73, and 0.82) in the C, P, and U datasets (Figure 5E). As expected, the correlation between the true cell number of taxa were relatively low (r_Spearman_ = 0.63, 0.58, and 0.52 for the C, P, and U datasets, respectively) (Table S11). The consistency of the true protein biomass of taxa and metaExpertPro results at species level was relatively low (r_Spearman_ = 0.2, 0.27, and 0.35) in the C, P, and U dataset (Figure S6G, Table S11).

Taken together, we found that filtering the taxa with at least three to five peptides led to the highest F-score at genus and species levels, and metaExpertPro achieved high accuracy in both diversity and biomass at genus level. The relatively low accuracy at species level might be due to that we used the Unipept-based taxonomy annotation. As a peptide-centric taxonomic annotation software, Unipept depends on taxon-specific peptides to identify taxa. However, the number of taxon-specific peptide sequences steadily decreases from higher to lower taxonomic rankings, with a particularly large drop between genus and species levels^47^. In addition, there are some species or even genera in the metaproteomic samples do not present in the NCBI taxonomy database, such as *Burkholderia xenovorans*, *Nitrosomonas europaeae*, *Pseudomonas denitrificans*, *Pseudomonas pseudoalcaligenes* and *Burkholderia* (Figure 5E, Figure S6G, marked in gray), which leads to false negative taxa. Nevertheless, Unipept is still the preferred software for taxonomy annotation in the absence of matched metagenomic data according to the previous study^30^. Here, we showed that metaExpertPro integrated Unipept can achieve high accuracy in the relative abundance estimation of genera (Figure 5E, Table S11).

### Negligible effects of public gut microbial gene catalog databases on DIA-MS-based proteome measurements

Three types of protein databases were commonly used in gut microbiota metaproteomic studies, including well-annotated public gut microbial gene catalog databases (*e.g.*, integrated gene catalog (IGC) of human gut microbiome^45^, Unified Human Gastrointestinal Protein (UHGP) catalog^48^), protein sequences that predicted from metagenome data from matched samples, and the merged databases of above two types of databases. To evaluate the impacts of databases on the peptide identifications in spectral library generation, we compared the peptide numbers in the five spectral libraries based on IGC+^49^, UHGP-90 (90% protein identity), matched metagenomic protein catalog database (MG), and their merged databases (MG_IGC+ and MG_UHGP-90) using 90 min gradient DDA-MS acquisition on timsTOF Pro of the 62 human fecal samples mentioned above (Figure 6A). The data showed that the spectral library based on IGC+ database identified the most peptides (284,681), followed by MG_IGC+ database (273,779), MG_UHGP-90 (273,338), UHGP-90 (271,751) and MG (261,986) (Figure 6B, Table S12). More specifically, 57.0% (194,485) of the peptides were commonly identified by all the spectral libraries. The spectral library based on MG contained the most unique peptides (21,296) (Figure 6C, Table S12). The identification rate of IGC+ spectral library was significantly higher than that of the other four databases. The identification rates (average of 30.6–31.8%) based on the five databases were comparable to the MetaPro-IQ^12^ results searching against matched metagenome (average of 34%) and IGC (average of 33%) (Figure 6D, Table S13). Overall, we found that in the spectral library generation step of metaExpert Pro, public gene catalog databases outperformed the matched metagenome database in terms of peptide identification. A similar conclusion has been proposed by Zhang *et al.* using MetaPro-IQ^12^.

**Figure 6.**
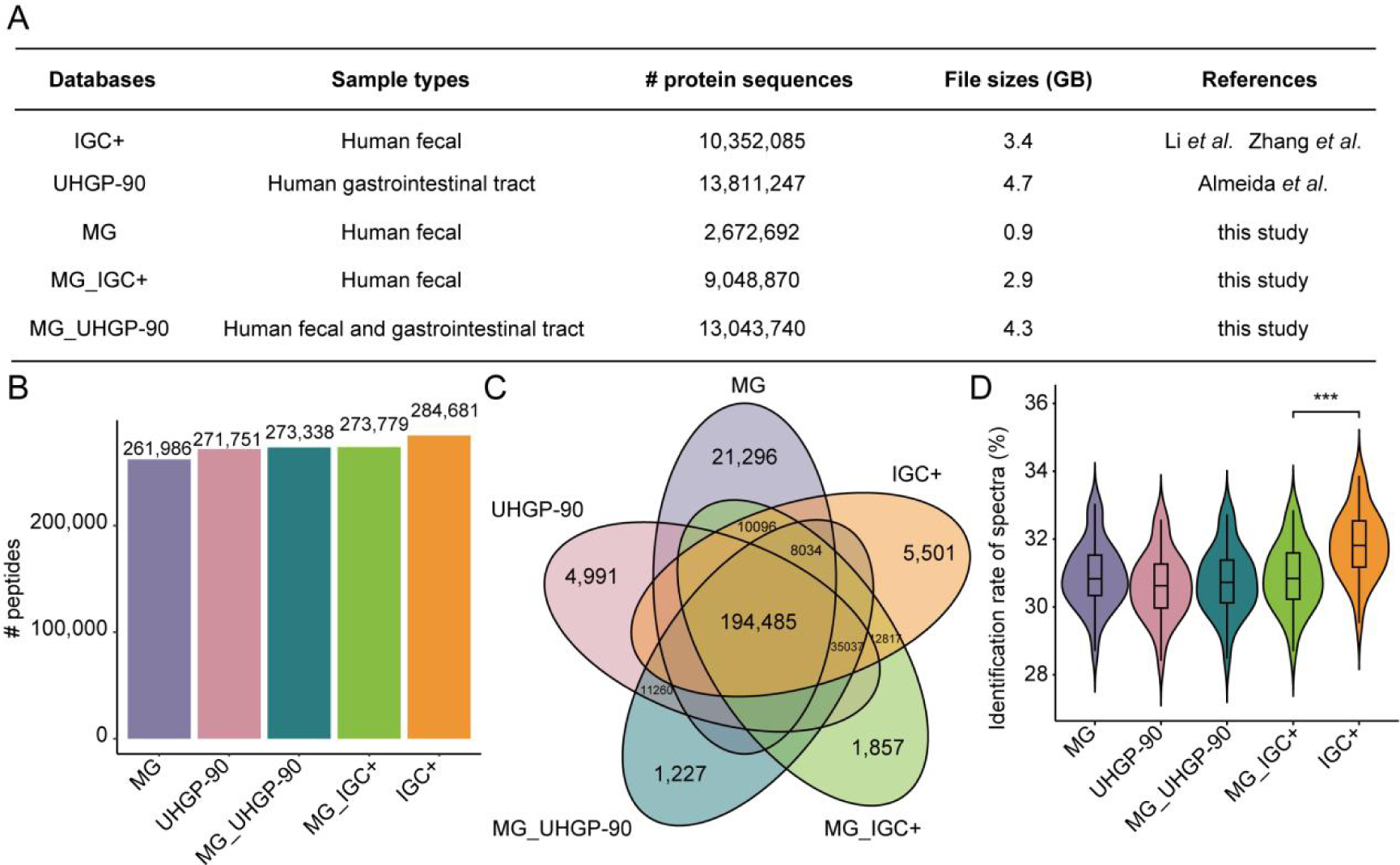
Comparison of the spectral libraries generated based on five different databases and DDA-MS data from human fecal samples. (A) The table lists the basic information of five protein databases, including IGC+, UHGP-90, MG, MG_IGC+, and MG_UHGP-90. The IGC+ database is the integrated gene catalog of human gut microbiome supplement with seven human gut fungal species, NCBI Virus, and gut microbial gene catalog of 28 mucosal-luminal interface samples. The UHGP-90 database is the Unified Human Gastrointestinal Protein catalog (UHGP-90) filtered by 90% protein identity. MG database is the matched metagenomic protein catalog database from 62 human feces. The MG_IGC+ and MG_UHGP-90 are the merged databases using MG and IGC+ or UHGP-90, respectively. The number of total peptides (B), the shared and unique peptides (C), and the identification rate of acquired MS spectra in each DDA-MS profile (D) in five spectral libraries were generated based on five databases and 30 ddaPASEF MS data (90 min gradient) from 62 human fecal samples. *p* value: * *p* < 0.05; ** *p* < 0.01; *** *p* < 0.001, **** *p* < 0.0001.

We further investigated the impacts of different public gene catalog databases on 60 min DIA-MS-based proteome measurements using two public gut microbial gene catalog databases (IGC+ and UHGP-90). High mapping ratios were obtained at COG (medium of 95.3% and 95.5%), KO (medium of 76.1% and 76.7%), and taxonomy (medium of 87.5% and 87.6%) levels with the two databases (Figure S7). The mapping ratio at the phylum level was comparable to the results of six human fecal data analyzed by glaDIAtor^21^ (∼70%). But the mapping ratio was less than that of glaDIAtor at the genus level (∼18% vs ∼40%), which may be because we used a stringent taxonomy filtering criterion of at least five peptides per taxonomy to ensure the accuracy of identification.

Next, we compared the richness per sample at eight levels and observed no significant differences between the two databases at all levels (Figure 7A, Table S14). At the peptide, COG, and KO levels, we also observed a high proportion of overlapped features (77–92%) between the two databases (Figure 7B). 84% of the genera and 86% of the species were identified by both databases, showing a high degree of consistency. The taxonomic and functional profiles identified by the two databases were also highly similar (Figure 7C, Table S15). In detail, at the taxonomic level, most of the peptides (99.4%) were assigned to the four major phyla of human gut microorganisms characterized by metagenomic data^50–53^, namely Bacillota (∼60%), Bacteroidota (∼30%), Actinomycetota (∼9%), and Pseudomonadota (∼1%). Also, the profiles of taxa were highly similar to that obtained by glaDIAtor (∼60% Bacillota, ∼10% Bacteroidetes, ∼7% Actinomycetota, and ∼0.5% Pseudomonadota). At the functional level, the largest functional categories included G ‘carbohydrate metabolism’ (∼18%), J ‘translation’ (∼16%), and C ‘energy metabolism’ (∼10%), which was in line with previous studies of human fecal metaproteomes^21,54^ (Figure 7C, Table S15). The abundance of human protein groups, microbial functions, and taxa also showed high correlation (medium of pairwise Spearman correlation coefficients = 0.95–0.97) between the two databases (Figure 7D, Table S16). Taken together, these results suggest the negligible effects of public gut microbial gene catalog databases on DIA-MS-based quantification at peptide, functional, or taxonomic levels. Therefore, matched metagenomic sequencing may not be required for the metaExpertPro and the results generated by metaExpertPro based on public databases could be directly comparable.

**Figure 7.**
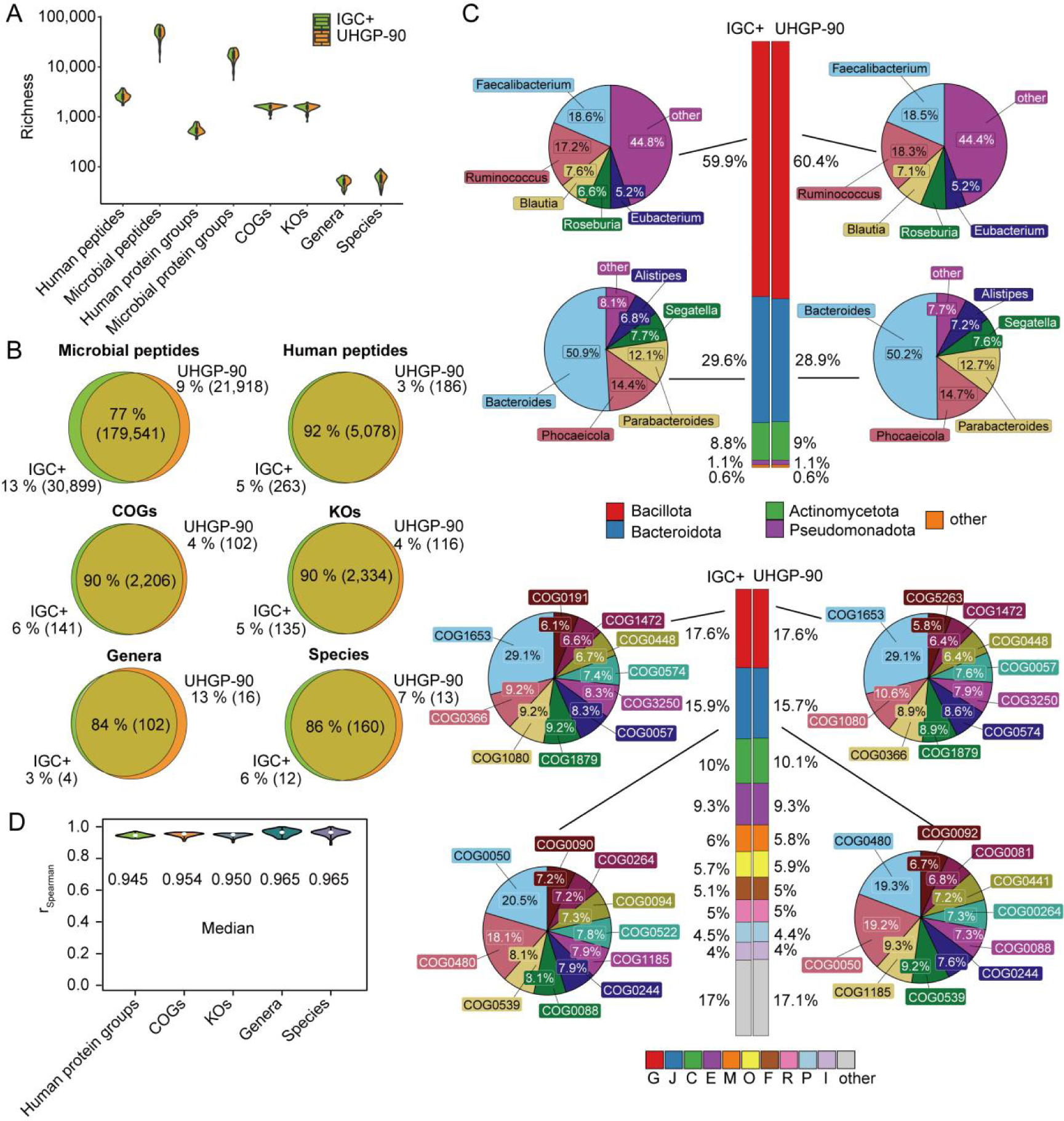
Negligible effects of public gut microbial gene catalog databases on DIA-MS based quantification. The analyses were based on 62 diaPASEF MS runs (60 min gradient) of 62 human fecal samples. (A) The number of quantitative peptides, protein groups, functions, and taxa per sample based on the IGC+ or UHGP-90 database. (B) The overlapped peptides, functions, and taxa in 62 human fecal samples based on IGC+ and UHGP-90 database. (C) The bar plots show the phylum-level taxonomic annotation of the peptides (upper) or COG-category-level functional annotation of protein groups (lower). The pie plots show the genus-level taxonomic annotation of the peptides (upper), or COG-level functional annotation of protein groups (lower) based on the IGC+ or UHGP-90 database. (D) The abundance correlation of human protein groups, functions, and taxa based on the IGC+ and UHGP-90 database. *p* value: * *p* < 0.05; ** *p* < 0.01; *** *p* < 0.001, **** *p* < 0.0001.

### metaExpertPro analysis revealed the functions associated with dyslipidemia and the potential interactions between the microbiota and the host

Dyslipidemia (DLP) is a disorder in lipid metabolism characterized by high levels of LDL-cholesterol and/or triglycerides and low HDL-cholesterol levels, which is considered a high-risk factor for cardiovascular disease^55,56^. Gut microbiota has been proved to be highly associated with dyslipidemia and related diseases^57^. However, the real functions of the microbiota associated with DLP are still unclear. The 62 GNHS subjects mentioned above included 31 subjects without DLP and 31 subjects with DLP. Here, we performed metaproteomic analysis on the fecal samples from these subjects to characterize the changes of microbial taxa, functions, and human protein groups in DLP. In total, we quantified 55,573 microbial protein groups and 993 human protein groups. The microbial protein groups were annotated as 2,347 COGs and 2,469 KOs. The microbial peptides were annotated as 106 genera and 172 species. About 87–97% of the identified protein groups, functions, and taxa were present in both non-DLP and DLP groups (Figure 8A). Two of the six genera uniquely identified in the DLP group (*Olsenella*^58,59^ and *Cloacibacillus*^60^) have previously been reported to show a positive association with serum lipids or obesity in mice, as well as in metabolically unhealthy obese human individuals. Among the eight genera uniquely identified in the non-DLP group, three have been reported to exhibit a negative association with DLP and obesity in mice. *Enterococcus*, a well-known probiotic, has been shown to alleviate obesity-associated dyslipidemia in mice^61,62^. *Lactococcus*, a potential antihyperlipidemic probiotic^63^, is also linked to insulin resistance and systemic inflammation, exerting an antiobesity effect^64^. *Turicibacter* is markedly reduced in mice fed with high-fat diet (HFD)^65^. A total of 56 COGs, 3 species, and 18 human proteins were significantly associated with DLP using General Linear Model (GLM) (*p*-value < 0.05 and | beta coefficient | > 0.2) (Figure S8 A–C, Table S17). The t-distributed stochastic neighbor embedding (t-SNE) analysis showed two close clusters corresponding to the DLP and non-DLP groups based on the associated microbial COGs, human proteins, and species, respectively (Figure 8 B–C, Figure S8D, Table S18). Wilcoxon Rank Sum Test was used to further verify the associations. The data showed that 34 of the associated microbial COGs were significantly differentially expressed between the two groups (Wilcoxon Rank Sum Test, *p* < 0.05) (Table S19). Functions related to the “Energy production and conversion” (two COGs in category C), “Lipid transport and metabolism” (two COGs in category I), “Transcription” (two COGs in category K), “Replication, recombination and repair” (three COGs in category L), and “Intracellular trafficking, secretion, and vesicular transport” (one COG in category U) showed significantly increased in DLP group. While the functions related to “Amino acid transport and metabolism” (two COGs in category E), “Lipid transport and metabolism” (one COG in category I), “Inorganic ion transport and metabolism” (two COGs in category P), “intracellular trafficking, secretion, and vesicular transport” (one COG in category U), and “defense mechanisms” (one COG in category V) showed significantly decreased in the DLP group (Figure 8D, Table S19). The results indicated an enhancement in energy production, conversion, lipid transport, and metabolism functionality in the gut microbiota of DLP patients. The increase of the functions in DNA repair pathways such as uracil-DNA glycosylase (UDG) functions was consistent with the metaproteomic results in pediatric IBD patients^49^. Defects in human amino acid transporters are linked to inherited metabolic disorders^66^. In this study, we observed a reduction in amino acid transport and metabolism within the human gut microbiota. This finding suggests potential drug targets that could be focused on microbial proteins related to amino acid transport. We also found that the functions related to bacteria-secreted protein toxins such as biopolymer transport protein ExbD and WXG100 family proteins YukE and EsxA were downregulated in the DLP group (Figure 8D, Table S19). Two species including *Blautia luti* and *Fusobacterium mortiferum* were significantly differentially altered in DLP (Figure S8E). Both species or their corresponding genera have been reported to be associated with metabolic disorders including obesity^67^, type 2 diabetes or hypercholesterolemia^68^.

**Figure 8.**
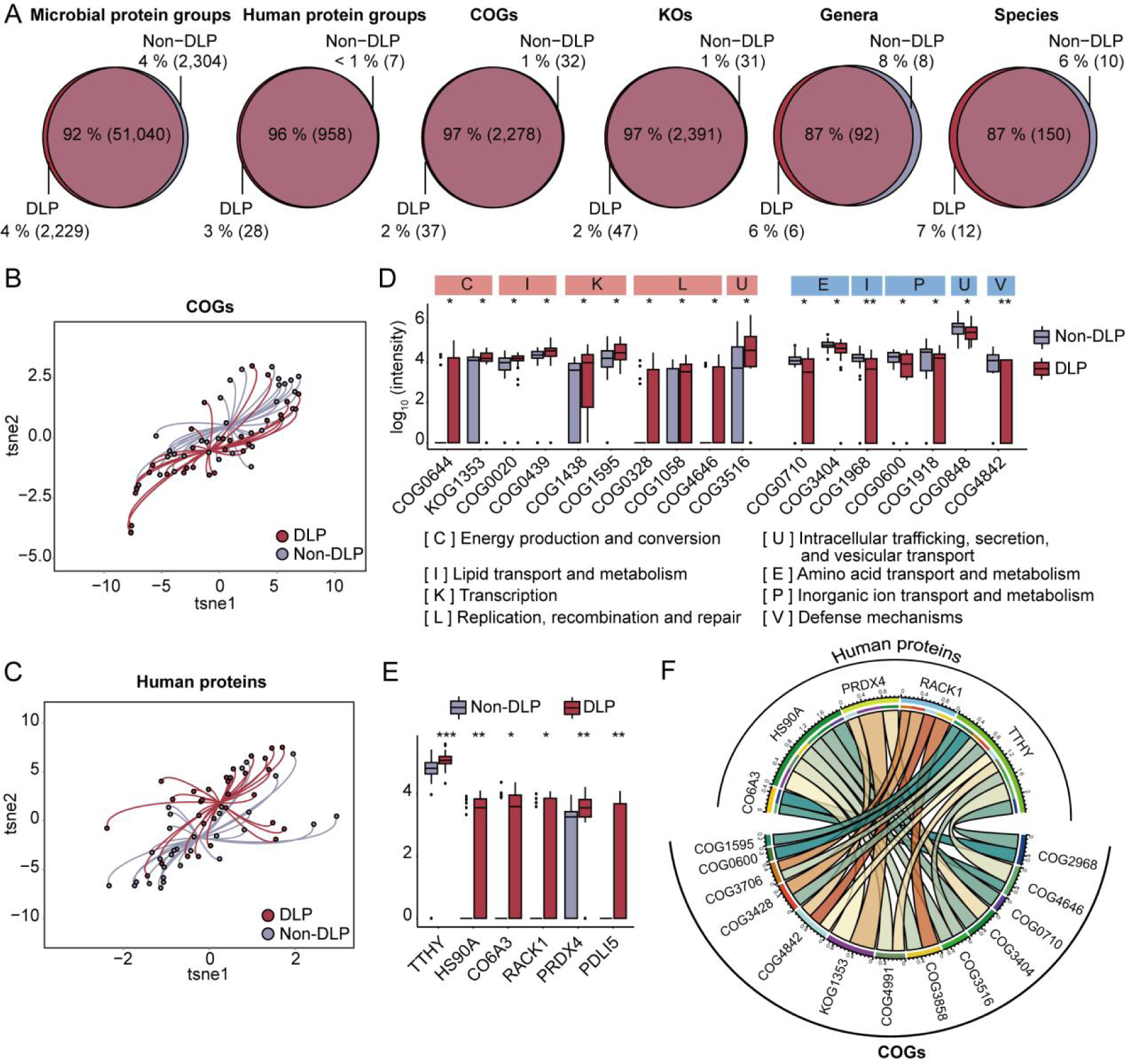
Proteins, functions, and taxa associated with DLP based on metaExpertPro workflow. The analyses are based on 62 diaPASEF MS runs (90 min gradient) of 62 human fecal samples collected from 31 non-DLP and 31 DLP subjects. (A) The overlapped quantitative proteins, functions, and taxa in DLP and non-DLP groups. The number of each section is labeled in the parenthesis. (B–C) The t-distributed stochastic neighbor embedding (t-SNE) visualization of DLP and non-DLP individuals calculated by significantly associated COGs (B) or human proteins (C) with DLP (General Linear Model (GLM) adjust the confounders of sex, age, and Bristol Stool Scale, *p*-value < 0.05 and | beta coefficient | > 0.2). (D) The intensity (log_10_ transformed) of significantly differentially expressed COGs (Wilcoxon Rank Sum Test, *p* < 0.05) belonging to the increased COG categories (red shadow) or the decreased categories (blue shadow). The COG categories are marked on the top of each COG. (E) The intensity (log_10_ transformed) of significantly differentially expressed human proteins between DLP and non-DLP groups (Wilcoxon Rank Sum Test, *p* < 0.05). (F) The co-expressed network between significantly changed COGs and human proteins in DLP. The co-expression between COGs and human proteins was determined by the Spearman correlation of their intensity in the 62 human fecal samples (| r_Spearman_ | ≥ 0.2, Benjamini-Hochberg [B-H] adjusted *p*-value <0.05). * *p* value < 0.05, ** *p* value < 0.01, *** *p* value < 0.001, **** *p* value < 0.0001.

One benefit of metaproteomic analysis was exploring the interactions between the host proteins and microbiota. Thus, we analyzed differentially expressed human proteins between the DLP and non-DLP groups using Wilcoxon Rank Sum Test. We identified six significant differentially expressed human proteins (*p* < 0.05). Interestingly, all of them were upregulated in the DLP group (Figure 8E, Table S19). Four human proteins including transthyretin (TTR), heat shock protein HSP 90-alpha (HS90A), small ribosomal subunit protein (RACK1), and peroxiredoxin-4 (PRDX4) have been reported to be related to obesity, diabetes, and hyperlipidemia based on serum or tissue samples^69–72^. However, it has not been reported that the dysregulation of these human proteins in human feces is also associated with dyslipidemia.

Next, we analyzed the co-expression between the six human proteins and the 34 differentially expressed COGs. With a threshold of | r_Spearman_ | ≥ 0.2 and Benjamini-Hochberg (B-H) adjusted *p*-value < 0.05, we screened out 25 co-expressed proteins and COGs (Figure 8F, Table S20). The human protein transthyretin (TTR) exhibited the strongest correlation with microbial COGs. Four positively correlated COGs were COG1595 (related to transcription), COG2968 (protein YggE), COG3516 (component TssA of the type VI protein secretion system), and COG4646 (adenine-specific DNA methylase). The other five negatively correlated COGs were COG0600 (ABC-type nitrate/sulfonate/bicarbonate transport system), COG3428 (membrane protein YdbT), COG3706 (Two-component response regulator, PleD family), COG4842 (secreted virulence factor YukE/EsxA, WXG100 family), and COG4991 (uncharacterized conserved protein YraI). Notably, the microbial function COG4842, a secreted virulence factor YukE/EsxA of the WXG100 family, exhibited negative correlations with three up-regulated human proteins (PRDX4, RACK1, and TTHY), indicating its significant role in the interaction with human proteins in the context of DLP. Taken together, the metaExpertPro-based metaproteomic analysis on DLP patients uncovered the alterations of microbial functions in DLP and the potential interactions between the microbiota and the host.

## Discussion

Due to the complexity of the samples, metaproteomic data analysis has inherent limitations of high dependency on databases, low efficiency of peptide identification rate (ID rate), the relatively low resolution of taxonomic identification, and large computer memory consumption. In this study, to solve the problems of low-efficiency ID rate and memory consumption, we used a library-based database search strategy in metaExpertPro, therefore our approach cannot eliminate the database dependency. FDR control poses another challenge in metaproteomics analysis due to large number of homologous bacterial sequences in the databases. In this study, benchmark tests using HeLa cell and bacteria mixture samples showed a low factual FDR (<5%). However, as the sample size and unmatched protein sequences in the database increase, controlling the factual protein group FDR becomes more challenging. Therefore, there is still a need for algorithms that can efficiently distinguish true positive spectra from highly similar spectra and employ stricter FDR filtering methods to ensure more accurate identifications.

Although our data showed negligible effects on the metaproteomic results based on two public gut microbial gene catalog databases and 62 human fecal samples, one cannot assume similar results can also be obtained with other gene catalog databases or other types of metaproteomic samples, such as soil microbiota and marine microbiota. Moreover, the Unipept-based taxonomic annotation still limits the resolution of accurate taxonomy identification at the species level due to the limited number of taxonomy-unique peptides. If matched metagenomic data is available, integrating metagenomic taxonomic information with Unipept has the potential to increase the number of taxonomy-unique peptides. This integration limits the potential species to those specific to the samples, leading to a higher count of taxonomy-unique peptides compared to considering all species from the NCBI taxonomy database. Thus, a novel taxonomic annotation software integrating metagenomic taxonomic information and Unipept has the potential to enhance the resolution of accurate taxonomy identification. Additionally, it is important to note that we did not observe any significantly associated microbial taxa, functions, or human proteins after correcting for multiple testing. This can be attributed to the limited number of samples used in our study, which consisted of 31 samples from individuals with dyslipidemia (DLP) and 31 samples from individuals without dyslipidemia (non-DLP). In order to obtain more accurate and reliable results, a larger sample size is required for future studies. Finally, this study and most published metaproteomic studies only focus on the proteins expressed by the host and microbiota; however, the proteins from foods and the environment may also play important roles in the hosts’ health and the metabolisms of microbiota. Therefore, despite these research advances, there is still much to discover in the metaproteome of the human gut.

### Conclusions

The metaExpertPro workflow provides a computational pipeline for metaproteomic analysis and shows a high degree of accuracy, reproducibility, and proteome coverage in the quantification of peptides, protein groups, functions, and taxa in human gut microbiota. The workflow is established by integrating the high-performance proteomic analysis tools and stringent filter criteria to ensure both in-depth and high accuracy measurments. The negligible effects of databases on the measurement of peptides, functions, and taxa indicate that matched metagenomic databases are not indispensable for metaExpertPro-based metaproteomic analysis, thus enabling direct comparison of metaproteomic data generated by metaExpertPro based on different public databases.

## Methods

### Human fecal sample collection

A total of 62 fecal samples were collected from 31 subjects without DLP and 31 subjects with DLP (40–75 years old) from the Guangzhou Nutrition and Health Study (GNHS)^39^. These individuals had not received any antibiotic treatment in the two weeks before biomaterial collections to avoid the effects of the antibiotic on the gut microbiome. The fecal samples were immediately homogenized, stored on ice, and then transferred to -80 ℃ within 4 h. Additionally, the corresponding metadata variables including age, gender, blood triglycerides (TG), total cholesterol (TC), low-density lipoprotein cholesterol (LDL), and high-density lipoprotein cholesterol (HDL) were also collected either by questionnaire or blood biochemical measurement. Dyslipidemia (DLP) was defined as one or more of the TG, TC, LDL, and HDL were abnormal or medical treatment for DLP^73^.

### Metaproteomic protein extraction and trypsin digestion

The gut microbiota was first enriched using differential centrifugation^74^. In detail, about 200 mg of feces were resuspended in 500 μL cold phosphate buffer (PBS) and centrifuged at 500 × g, 4°C for 5 min. Then the supernatant was transferred into a new tube. The above process was repeated three times. All the supernatants were combined (about 1.5 mL) and centrifuged at 500 × g, 4°C for 10 min to remove the debris in the fecal samples. Then, the microbial cells were collected by centrifugation at 18,000 × g, 4°C for 20 min. Next, the microbial pellets were used for protein extraction^75^. Briefly, 250 μL lysis buffer (4% w/v SDS and cOmplete Tablets (Roche) in 50 mM Tris-HCl, pH = 8.0) was added into the microbial pellets and the mixture was boiled at 95 °C for 10 min. Then, the mixture was ultrasonicated at 40 Khz (SCIENTZ) for 1 h on ice. Finally, to further discard the cell debris, the mixture was centrifuged at 18,000 × g for 5 min, and the proteins were precipitated overnight at -20 °C using a 5-fold volume of acetone. Next, the in-solution digestion method^76,77^ was performed as follows. After purifying (washing by acetone) and re-dissolving (using 8 mM urea and 100 mM ammonium bicarbonate) the precipitated proteins, about 50 μg proteins from each sample were reduced with 10 mM tris (2-carboxyethyl) phosphine (TCEP, Adamas-beta) and then alkylated with 40 mM iodoacetamide (IAA, Sigma-Aldrich). Proteins were pre-digested with 0.5 μg trypsin (Hualishi Tech) for 4 h at 32 °C. Then the proteins were further digested with another 0.5 μg trypsin for 16 h at 32 °C. The tryptic peptides were desalted using solid-phase extraction plates (ThermoFisher Scientific, SOLAµ™) and then freeze-dried for storage. Dried peptides were finally resuspended in a solution (2% acetonitrile, 98% water, and 0.1% formic acid [FA]) before MS acquisition.

### Metagenomic DNA extraction, sequencing, and gene prediction

The metagenomic raw data was derived from the previous study^78^. Briefly, the raw sequencing reads were first filtered and trimmed with PRINSEQ (version 0.20.4)^79^ for quality control. The raw reads aligned to the human genome (*H. sapiens*, UCSC hg19) were removed using Bowtie2 (version 2.2.5)^80^. Then, the remaining reads were used for metagenomic assembly using MEGAHIT (version 1.2.9)^81^ and binning the contigs with MetaBAT (version 2.12.1)^82^ by default parameters. We further clustered and de-replicated the Metagenome-Assembled Genome (MAGs) at an estimated species level (ANI ≥ 95%) using dRep (version 3.0.0)^83^. The minimum genome completeness and maximum genome contamination were set to 75 and 25, respectively. Protein-coding sequences (CDS) for each MAG were predicted and annotated with Prokka (version 1.13.3)^84^. All the predicted protein sequences were compiled to generate the MG protein database. Cd-hit (version 4.8.1)^85^ was used for the integration of MG and IGC+^49^ or UHGP^48^ database with the following parameters: -c 0.95 -n 5 -M 16000 -d 0 -T 32.

### High-pH reversed-phase fractionation

For the 62 fecal samples, approximately 5 μg peptides were collected from each tryptic peptide sample to form a pooled sample for high-pH fractionation. The pooled sample was then fractionated using high-pH reversed-phase liquid chromatography (LC). The mobile phase of buffer A was water with 0.6% ammonia (pH = 10), and buffer B was 98% acetonitrile and 0.6% ammonia (pH = 10). Specially, about 300 μg tryptic peptides were separated using a nanoflow DIONEX Ultimate 3000 RSLC nano System (ThermoFisher Scientific) with an XBridge Peptide BEH C18 column (300 Å, 5 μm× 4.6 mm × 250 mm) at 45 °C. A 60 min gradient from 5% to 35% buffer B with a flow rate of 1 mL/min was applied. A total of 60 fractions were collected and further combined into 30 fractions. Finally, the fraction samples were freeze-dried and re-dissolved in 2% acetonitrile with 98% water and 0.1% FA.

### DDA mass spectrometry acquisition for library generation

The fractionated peptides were first spiked with iRT (Biognosys)^86^. For the timsTOF Pro (Bruker) based DDA mass spectrometry acquisition, two gradients of 90 min and 60 min were used, respectively. The 90 min LC gradient was linearly increased from 2% to 22% buffer B for 80 min, followed by a second linear gradient from 22% to 35% buffer B for 10 min (buffer A: 0.1% FA in water; buffer B: 0.1% FA in ACN). The 60 min LC gradient was linearly increased from 5% to 27% buffer B for 50 min, followed by a second linear gradient from 27% to 40% buffer B for 10 min. The peptides were loaded at 217.5 bar on a precolumn (5 μm, 100 Å, 5 mm × 300 μm I.D.) in 0.1 % FA/water and then separated by a nanoElute UHPLC System (Bruker Daltonics) equipped with an in-house packed 15 cm analytical column (75 μm ID, 1.9 μm 120 Å C18 beads) at a flow rate of 300 nL/min. The timsTOF Pro was operated in ddaPASEF mode with 10 consecutive PASEF MS/MS scans after a full scan in a total cycle. The capillary voltage was set to 1400 V. The MS and MS/MS spectra were acquired from 100 to 1700 m/z. The TIMS section was operated with a 100 ms ramp time and a scan range of 0.6–1.6 V·s/cm^2^. A polygon filter was used to filter out singly charged ions. For all experiments, the quadrupole isolation width was set to 2 Th for m/z < 700 and 3 Th for m/z > 800. The collision energy was ramped linearly as a function of mobility from 20 eV at 1/K_0_ = 0.6 V·s/cm^2^ to 59 eV at 1/K_0_ = 1.60 V·s/cm^2^.

For the Orbitrap Exploris™ 480 mass spectrometer (ThermoFisher Scientific Inc.) based DDA mass spectrometry acquisition, the fractionated peptides spiked with iRT were loaded onto a pre-column (3 μm, 100 Å, 20 mm × 75 mm i.d., Thermo Fisher Scientific, USA) using a Thermo Scientific UltiMateTM 3000 RSLCnano LC a U3000 LC system. The peptides were then separated at a flow rate of 300 nL/min using a 60 min LC gradient on an in-house packed 15 cm analytical column (75 μm ID, 1.9 μm, C18 beads) with a linear gradient from 5% to 28% buffer B for 60 min. Next, the column was washed with 80% buffer B. The mobile phase B consisted of 0.1% formic acid in MS-grade ACN, while the mobile phase A consisted of 0.1% formic acid in 2% ACN and 98% MS-grade water. The eluted peptides were analyzed by an Exploris 480 MS with the FAIMS Pro (High field asymmetric waveform ion mobility spectrometry) interfacing in standard Data-dependent acquisition (DDA) acquisition mode. Compensation voltage was set at two different CVs, -42 and -62 V, respectively. Gas flow was applied with 4 L/min with a spray voltage set to 2.1 kV. The DDA was performed using the following parameters. MS1 resolution was set at 60,000 at m/z 200 with a normalized AGC target of 300%, and the maximum injection time was set to 20 ms. The scan range of MS1 ranged from 350–1200 m/z. For MS2, the resolution was set to 15,000 with a normalized AGC target of 200%. The maximum injection time was set as 20 ms for MS1. Dynamic exclusion was set at 30 s. Mass tolerance of ± 10 ppm was allowed, and the precursor intensity threshold was set at 2e4. The cycle time was 1 second, and the top-abundance precursors (charge state 2−6) within an isolation window of 1.6 m/z were considered for MS/MS analysis. For precursor fragmentation in HCD mode, a normalized collision energy of 30% was used. All data were acquired in centroid mode using positive polarity and peptide match and isotope exclusion were turned on.

We obtained a total of 90 DDA-MS raw data profiles. These included 30 profiles from timsTOF Pro MS instrument with a 60 min gradient, 30 profiles from timsTOF Pro MS instrument with a 90 min gradient, and 30 profiles from Exploris 480 MS instrument with a 60 min gradient.

### DIA mass spectrometry acquisition for peptide and protein quantification

For the timsTOF Pro-based DIA-MS acquisition, 300 ng peptides were trapped at 217.5 bar on the precolumn and then separated along the 60 min LC gradient same as the ddaPASEF LC gradient mentioned above. The ion mobility range was limited to 0.7–1.3 V·s/cm^2^. Four precursor isolation windows were applied to each 100 ms diaPASEF scan. Fourteen of these scans covered the doubly and triply charged peptides’ diagonal scan line in the m/z ion mobility plane. The precursor mass range 384–1087 m/z was covered by 28 m/z narrow windows with a 3 m/z overlap between adjacent ones. Other parameters were the same as the setting in the ddaPASEF acquisition.

For the Exploris 480-based DIA-MS acquisition, 500 ng peptides were separated by the LC methods with a slight modification from the DDA-MS LC methods. The initial phase B of the gradient was increased from 5% to 7% to get a more effective time for separation. The Spray voltage of FAIMS was set to 2.2 kV. The other FAIMS settings were consistent with those of the DDA-MS acquisition. In DIA mode, full MS resolutions were set to 60,000 at m/z 200 and the full MS AGC target was 300% with an IT to 50 ms. The mass range was set to 390–1010. The AGC target value for fragment spectra was set at 2000%. 15 isolation windows of 15 Da were used for -62V compensation voltage with an overlapped of 1 Da, and 19 isolation windows of 20 Da were used for -42V compensation voltage with an overlapped of 1 Da. The resolution was set to 15,000 and the IT to 54 ms. The normalized collision energy was set at 32%.

Overall, 62 diaPASEF raw data profiles and 60 DIA-MS (Exploris 480) raw data profiles were obtained for the human fecal samples.

### Comparison of metaExpertPro with other metaproteomics software tools

We firstly incorporated the comparison of DDA-MS-based peptide identifications among ProteoStorm^16^, metaLab^13^, glaDIAtor^42^, and metaExpertPro using the same raw data, database, and parameters. Specially, six DDA-MS files of human fecal samples from dataset PXD008738^42^ were searched against the integrated gene catalog (IGC) database using ProteoStorm, glaDIAtor, and metaExpertPro, respectively. Enzyme specificity was set to “Trypsin/P” with maximum one missed cleavage. Precursor mass tolerance and fragment mass tolerance were set at 10 ppm and 0.02 Da, respectively. All the tests were performed on a computer with AMD EPYC hardware and 512GB RAM.

### Multivariate statistical analysis

The intensity values at peptide, protein, functional and taxonomic levels were log_10_ transformed for statistical analysis. The reproducibility of the quantitative proteins, functions, and taxa in biological replicate samples was estimated by Spearman correlation. The intensity comparisons of the identified peptides and protein groups between glaDIAtor^42^ and metaExpertPro were conducted using Wilcoxon Rank Sum Test. The COGs, KOs, human proteins, and species significantly associated with DLP were determined by General Linear Model (GLM)^87^ (adjust the confounders of sex, age, and Bristol Stool Scale, *p*-value < 0.05 and | beta coefficient | > 0.2). The differentially expressed human proteins, COGs, and species were identified by Wilcoxon Rank Sum Test (*p*-value < 0.05). t-SNE was performed using the Rtsne package (version 4.1.3). The co-expressed COGs and human proteins were identified using the Spearman correlation of their abundance in 62 human fecal samples (| r_Spearman_ | ≥ 0.2, Benjamini-Hochberg [B-H] adjusted *p*-value <0.05).

## Declarations

### Ethics approval and Consent to participate

The study protocols of the Guangzhou Nutrition and Health Study were approved by the Ethics Committee of the School of Public Health at Sun Yat-sen University and the Ethics Committee of Westlake University. Written informed consent was obtained from all participants.

### Consent for publication

Not applicable.

### Authors’ contributions

T.G., J.Z, and Y.C. designed and supervised the project. Z.M., L.Z., and H.Z. collected the samples and metadata. Y.S., Z.X., S.L., W.J., H.G., Y.X., L.Y., and X.C. generated the data. Y.S., Z.X., S.L., H.Z., and Y.Z. analyzed the data. Y.S., Z.X, S.L., and T.G. drafted the manuscript with inputs from all co-authors. Y.S., Z.X., and S.L. contributed equally to this work.

## Acknowledgements

We thank Dr. Fengchao Yu from the Alexey I. Nesvizhskii group at University of Michigan for the assistance in establishing the pipeline. We thank Dr. Xu Zhang from the Daniel Figeys group at University of Ottawa for generously providing the human gut gene catalog database, IGC+. We thank Laura L. Elo group at University of Turku and Åbo Akademi University for their support in the processing of glaDIAtor. We thank Dr. Guojie Cui for the discussion. We thank Westlake University Supercomputer Center for assistance in data generation and storage.

